# Role of *Staphylococcus aureus*’s Buoyant Density in the Development of Biofilm Associated Antibiotic Resistance

**DOI:** 10.1101/2023.12.23.573193

**Authors:** Sarah Kispert, Madison Liguori, Cody Valikaneye, Chong Qiu, Shue Wang, Nan Zhang, Huan Gu

**Author notes:** Equal contribution: Those two authors contributed equality to this study (Sarah Kispert, Madison Liguori). Corresponding author, Huan Gu.

## Abstract

Bacterial biofilms are clusters of bacterial cells that form at various interfaces, including those between air and liquid or liquid and solid. Due to their roles in enhancing wastewater treatment processes, and their unfortunate propensity to cause persistent human infections through high antibiotic resistance, understanding and managing bacterial biofilms is of paramount importance. A pivotal stage in biofilm development is the initial bacterial attachment to these interfaces. However, the determinants of bacterial cell choice in colonizing an interface first and heterogeneity in bacterial adhesion remain elusive. Recent research has unveiled variations in the buoyant density of *Staphylococcus aureus* cells, irrespective of their growth phase. Cells with a low cell buoyant density, characterized by fewer cell contents, exhibited greater resistance to antibiotic treatments (100 μg/mL vancomycin) and favored biofilm formation at air-liquid interfaces. In contrast, cells with higher buoyant cell density, which have richer cell contents, were more vulnerable to antibiotics and predominantly formed biofilms on liquid-solid interfaces when contained upright. In essence, *S. aureus* cells with higher buoyant cell density are more inclined to adhere to upright substrates.

## 1. Introduction

Biofilms consist of bacteria that adhere to surfaces and are enveloped in a self-secreted extracellular polymeric substance (EPS)^1–3^. Their robust resistance to antibiotics arises from the protective shield offered by the EPS and the presence of dormant cells, termed “persisters”^4, 5^. As per data from the Centers for Disease Control (CDC, 2007), bacterial biofilms were linked to nearly 1.7 million hospital-related infections, translating to medical expenses of $11 billion^6^. The annual economic burden of infections related to biofilms in the U.S. is approximately $94 billion, contributing to over half a million fatalities in the United States^7^. Given their profound impact, biofilm research and control have been rigorously pursued for the last thirty years. The initiation of biofilm formation is marked by bacterial attachment to solid substrates, like medical implant tissues^8^. The mechanisms governing the shift of bacteria from a free moving (or planktonic) phase to structured biofilms remain ambiguous, spurring the focus of our study. Insights from this study endeavor aim to enrich the formulation of potent strategies for biofilm prevention and eradication.

Bacterial buoyant density indicates a bacterial cell’s propensity to float or settle within a fluid medium like a growth solution^9–11^. When comparing planktonic cells of varying buoyant densities, those with a higher buoyant density tend to settle more rapidly due to gravitational forces. Notably, in 1984, Kubitschek et al. reported variations in the buoyant density of *Escherichia coli* cells, exhibiting a consistent normal distribution irrespective of their growth pace^10^. Recent findings suggest a correlation between bacterial buoyant density, growth rate, and cell age^12–14^. However, this might not hold for persisters, a subset of bacteria known for having fewer cell organelles than their typical counterparts ^15, 16^. Intriguingly, persisters are consistently present in almost all bacterial populations, independent of growth rates or population age. Given the observed relationship between organelle count and buoyant density^16^, we postulate that cells with elevated buoyant densities, due to their heightened metabolic activities, are more inclined to migrate towards and initially attach to upright substrates, driven by gravitational forces.

To validate our hypothesis, we utilized sucrose density gradient centrifugation to separate a *Staphylococcus aureus* (*S. aureus*) population based on varying buoyant cell densities. This method then helped us assess their biofilm formation capabilities and antibiotic susceptibility. Although sucrose density gradient centrifugation is commonly applied in separating nanoparticles^17, 18^, liposomes^19^, membranes from tissues and cells^20^, ribonucleic acids^21^, or viruses^22, 23^, its incorporation in bacterial studies remains limited. Our research endeavors to harness this established technique to discern the influence of bacterial buoyant density on biofilm development and related antibiotic resistance. In this study, we selected *S. aureus* as the model microorganism due to its clinical importance and its resemblance to micron-scale biological entities. *S. aureus* is known as an opportunistic pathogen that can cause multidrug-resistant afflictions ranging from skin and respiratory infections^24, 25^ to food poisoning^26^. These spherical bacteria, are devoid of external structures like flagella, and lack motility^27^. Consequently, *S. aureus* cells can be viewed as static biological particles with minimal interactions, enabling an exclusive focus on buoyant density.

## 2. Results

### 2.1 For any isogenic *S. aureus* population, there was a distribution of cell buoyant density, regardless of the population’s age

First, we conducted a gravity-driven sedimentation experiment using stationary phase *S. aureus* cells to quantitatively evaluate if there is a difference in the rate of sedimentation among the cells in an isogenic population. Using Stokes Law based on the equilibrium of forces, the sedimentation speed (*v*) can be calculated using Equation (1) below.

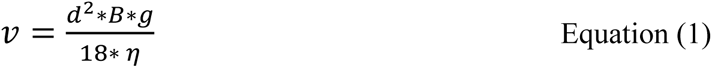

where *d* is particle/cell diameter; *B* is buoyant/particle-to-liquid relative density; *g* is gravitational constant; and *η* is the dynamic density of the mixture.

Based on Equ. (1), the sedimentation rate is proportional to buoyant density of cells. Hence, the faster sedimentation rate represents larger buoyant density.

To obtain an isogenic population, we used a single bacterial colony on an LB agar plate to inoculate bacterial growth. By carefully introducing a 15.5 hr-old isogenic *S. aureus* population at the top of 5 mL pre-chilled 0.85% NaCl solutions in 15 mL Falcon Tubes and letting the cells settle for 15 min on ice, the difference in the rate of sedimentation was evaluated. The 0.85% NaCl solutions were used in this experiment to avoid the effect of cell metabolic activities and division on the rate of sedimentation. The time of sediment was chosen as 15 min because it was the time that takes for chemical diffusion to reach equilibrium^28^. By quantifying the sedimentation rates of cells in *S. aureus* in an isogenic population (1.55 ± 0.74 × 10^9^ cells or 100%), we found that cells sink at different rates (Fig. 1). It is worth noting that there were always 0.41 ± 0.05% or 6.37 ± 0.79 × 10^8^ cells that did not sediment in 15 min. Although the percentage was small, the absolute number of these cells was significant among the total population. With the sedimentation rates in Fig. 1, 51.17 ± 6.24% cells got the closest to the bottom surface of the 15 mL Falcon Tube and were ready to make initial attachments. This portion of cells sediment faster, possibly due to the larger buoyant density.

**Figure 1.**
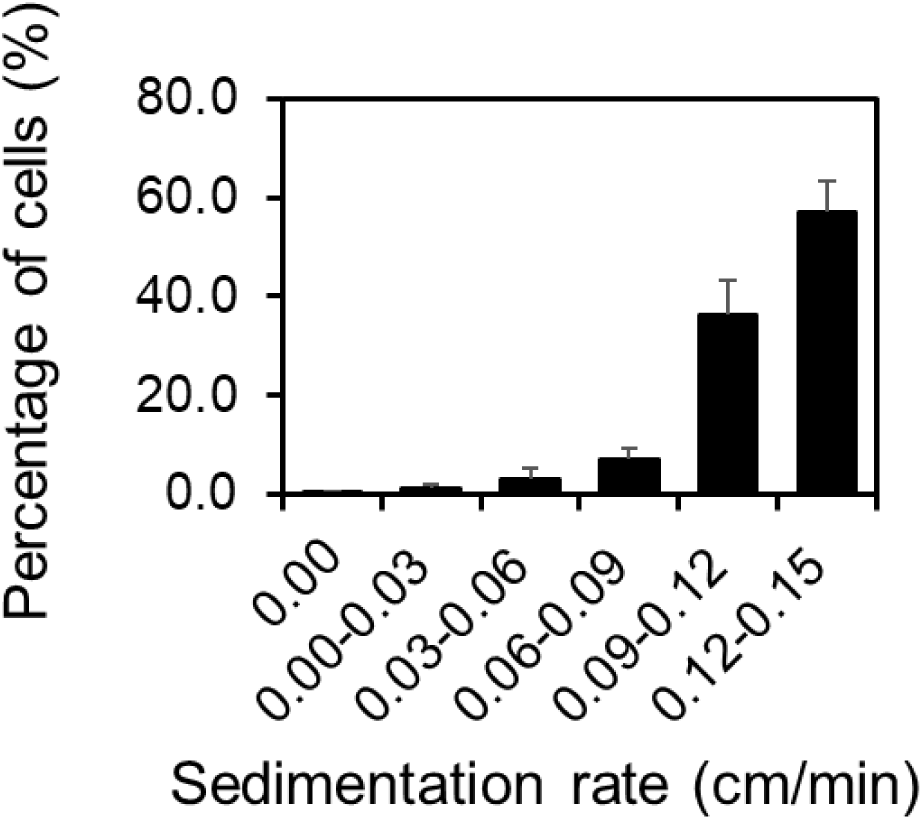
Gravity-driven sedimentation experiment. Percentage of *S. aureus* cells sediment at various speeds. In 15 mL Falcon tube with 5 mL 0.85% NaCl solution, cells sediment with 0.12-0.15 cm/min were the ones that were closest to the bottom of the upright 15 mL falcon tube and ready for initial attachment after 15 min of gravity-driven sedimentation. This experiment was repeated biologically four times (N = 4). Standard errors were shown.

### 2.2 Correlating sedimentation rates with cells’ buoyant density using sucrose density gradient centrifugation

The different sedimentation rates observed above indicate various buoyant density. To specifically characterize the buoyant density of cells in an isogenic *S. aureus* population, we used the sucrose gradient differential centrifugation. Because we used sucrose solutions with known densities (998, 1000, 1018, 1038, 1081, 1176, or 1230 kg/m^3^), cells stopped in the sucrose solution with the same density (or *v* = 0 when *B* = 0 kg/m^3^ based on Equ. 1). By pulling an isogenic *S. aureus* population through a stack of sucrose solutions (or sucrose density gradient centrifugation), cells were separated based on their density. Using this method, we found that more than 86.32% of a 15.5-hr old *S. aureus* population had a mass density higher than 1230 kg/m^3^ (Fig. 2a). This could lead to a buoyant density (B) of 234 kg/m^3^ if those cells are suspended in solutions with a mass density like water (998 kg/m^3^), leading to sedimentation in response to gravity. It is worth noting that there is a heterogeneity in *S. aureus* cells’ density and there is a subpopulation (0.0005 ± 0.0002%) with a mass density smaller than that of water. Although the size of this subpopulation is small, the absolute cell number is significant (4.34 ± 1.66 × 10^3^ among 8.25 ± 2.66 × 10^8^ cells).

**Figure 2.**
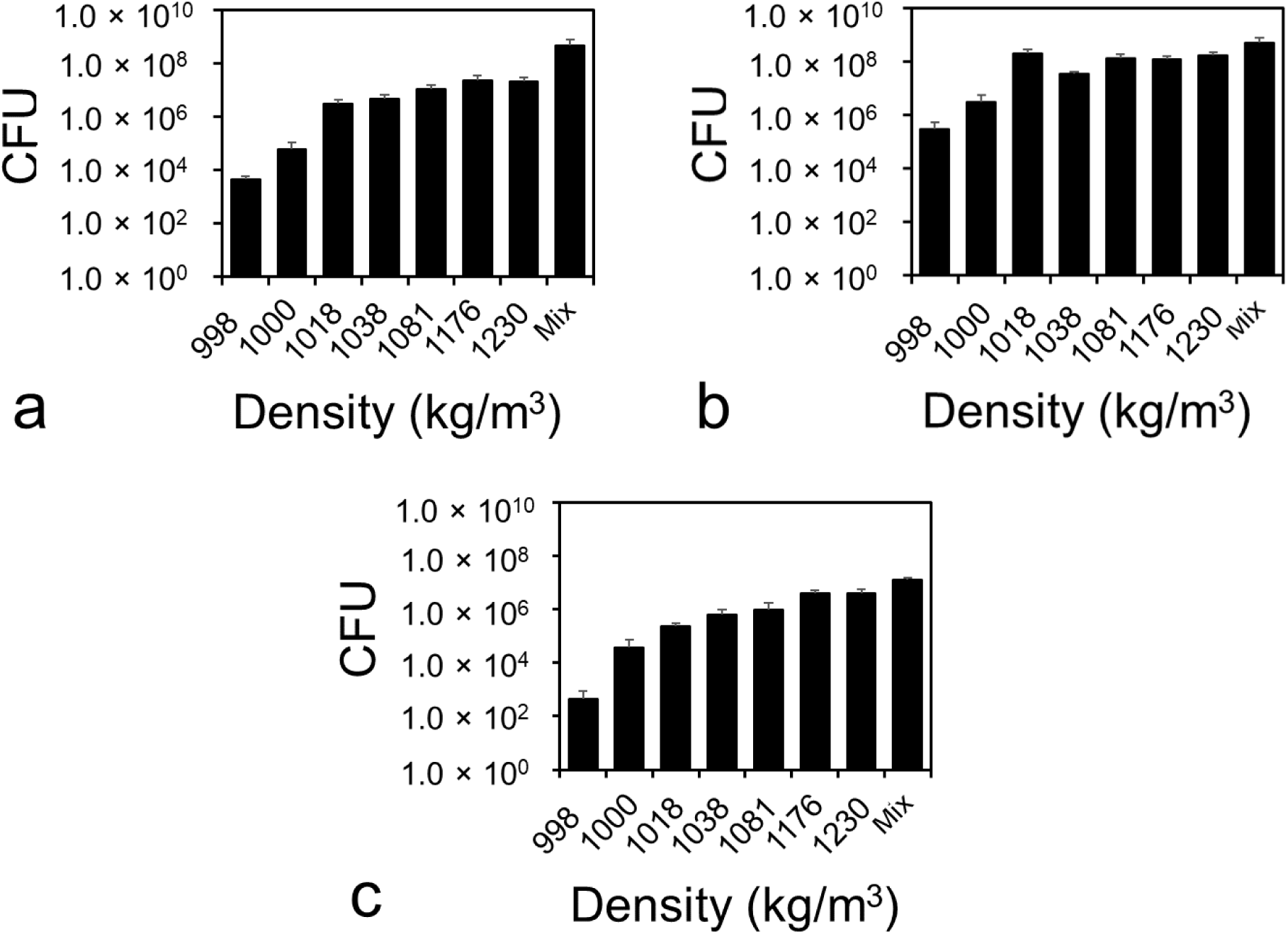
Sucrose gradient differential centrifugation. The distribution of the cells in a 15.5 (a), 18 (b), and 2 (c)-hr old *S. aureus* populations. The total population (mix) were pulled through a stack of sucrose solutions with various mass densities (998, 1000, 1018, 1038, 1081, 1176, or 1230 kg/m^3^) and separated based on their mass density. The absolute number of cells were quantified using the drop plate method and CFU^29^. Each separation was repeated five times biologically (N = 5). Standard errors are shown.

This heterogeneity in cell density is true regardless of growth age. However, the pattern of cell distribution at different densities varies according to growth age (Fig. 2b&c). For instance, in an 18-hr old *S. aureus* population, there were no *S. aureus* cells with densities higher than 1230 kg/m^3^ and the percentage of *S. aureus* cells with densities smaller than that of water increased to 0.06 ± 0.05% or 3.06 ± 2.30 × 10^5^ among 5.06 ± 2.24 × 10^8^ cells. It is worth noting that the distribution of 18-hr old *S. aureus* cells does not follow normal distribution. In a 2-hr old exponential *S. aureus* population, 23.7% cells had a mass density larger than 1230 kg/m^3^ and the percentage of *S. aureus* cells with a density smaller than that of water was 0.0035 ± 0.0031% or 4.63 ± 4.13 × 10^2^ among 1.33 ± 0.26 × 10^7^ cells. Although the percentage of cells with a density smaller than that of water in the exponential phase *S. aureus* population was higher than that in a 15.5-hr old *S. aureus* population, the absolute number of cells remained low.

### 2.3 Cells with larger density formed better biofilms compared to those with lower mass density

Using sucrose density gradient centrifugation, we separated 15.5 hr-old *S. aureus* cells based on their mass density and compared their biofilm formation capabilities on the bottom surface of 96-well plates. The amount of biofilms formed by cells with different cell densities was quantified using crystal violet assay^30^ (Fig. 3). As shown in Fig. 3, cells with higher mass density (1230 kg/m^3^) formed more biofilms on the bottom surface of 96-well plates compared to cells with lower mass density (998 or 1000 kg/m^3^), which supports our hypothesis (t-test, *p* = 0.028 and 0.026 <0.05). They also formed slight better biofilms compared to those formed by *S. aureus* cells without separation (t-test, *p* = 0.21 > 0.05). Since *S. aureus* cells without separation is a mixture of cells with both low and high densities, these results corroborate our hypothesis. Because we matched the sucrose background and cell numbers for inoculating biofilm growth, this difference in biofilm formation observed in Fig. 3 was solely due to the mass density of *S. aureus* cells.

**Figure 3.**
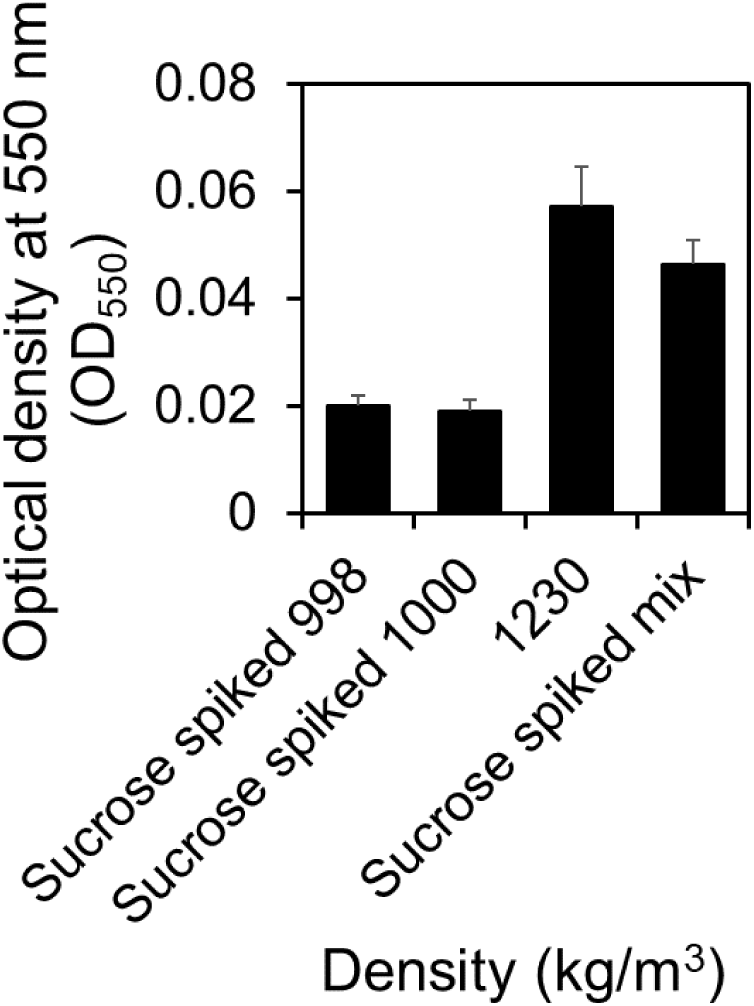
Biofilms formed by cells with different densities. The amount of biofilms formed by cells with low-level mass densities (998 and 1000 kg/m^3^) and high-level mass density (1230 kg/m^3^) after 48 hours of incubation. The biofilms formed by the mixture of cells with varying densities before separation were used as controls. The biofilm formation was repeated four times biologically (N = 4). Standard errors are shown.

### 2.4 Cells with larger density revived faster from nutrient deprivation than those with lower density when nutrients were supplemented

To investigate the correlations between biofilm formation and cell density, we evaluated the growth rate of cells with low-level mass densities (998 and 1000 kg/m^3^) and high-level mass density (1230 kg/m^3^) after they were resuspended in the solutions with abundant nutrients (Fig. 4).

**Figure 4.**
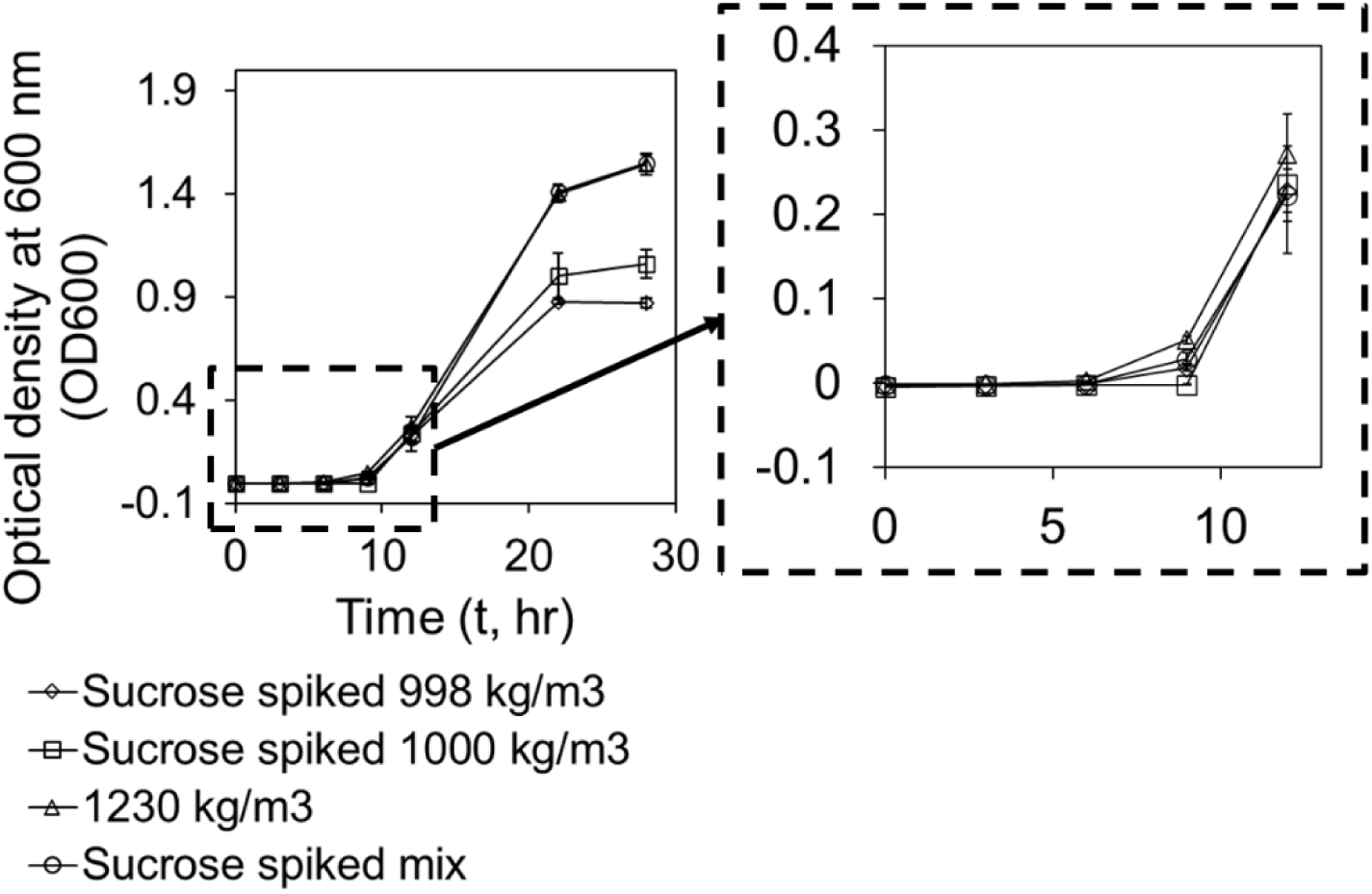
Cell growth rate. Cells with low-level mass densities (998 and 1000 kg/m^3^) and high-level mass density (1230 kg/m^3^) were isolated using sucrose density gradient centrifugation. Then, cells were resuspended into fresh LB with the same initial cell density. The optical cell density at 0, 3, 6, 9, 12, 22, and 28 h after inoculation was measured using spectrophotometer. Each measurement was repeated biologically three times (N =3). Standard errors are shown.

Based on the results, we found that *S. aureus* cells with high-level density (1230 kg/m^3^) started reproducing first at 6 h after inoculation (t-test, *p* = 0.024, 0.033, or 0.049 < 0.05 compared to sucrose spiked cells with densities of 998 and 1000 kg/m^3^ or sucrose spiked mix, respectively), indicating that they are different. At 12 h after inoculation, growth rate of cells with low-level densities caught up. However, at 22 h after inoculation, the growth rate of sucrose spiked cells with the density of 998 kg/m^3^ became significantly slower than cells with high-level density (t-test, *p* = 0.0003 and 0.0000 compared to cells with density of 1230 kg/m^3^ and sucrose spiked mix, respectively). At 28 h after inoculation, their growth rate was even slower than sucrose spiked cells with the density of 1000 kg/m^3^ (t-test, *p* = 0.057). Since we matched the sucrose concentration of cells with different densities after sucrose density gradient centrifugation, this difference in growth rate was solely due to the heterogeneity in cell density.

### 2.5 Higher percentage of cells with higher buoyant density had DNA

To investigate the mechanism behind the heterogeneity in cell density, we first evaluated the correlation between the DNA and mass density (Fig. 5). We used Acridine Orange (AO) to labeled the DNA^31^. Using different illumination methods (Brightfield vs. green fluorescence) to look at the cells in a 15.5-hr S. aureus population after separation, we found that the distribution of cells with DNA labeled with green fluorescence followed the similar distribution as the cells that can be quantified using CFU (DNA in Fig. 5). However, the total number of cells identified using Brightfield illumination were significantly more than that quantified using CFU, especially when the cell density was close to that of water (998 kg/m^3^). When the cells density was 998, 1000, 1018, and 1038 kg/m^3^, only 6.90, 0.62, 5.30, and 17.01% of the cells visualized by Brightfield had their DNA labeled with AO (t-test, *p* = 0.0006, 0.0541, 0.0283, and 0.0100, respectively). This percentage increased to 69.92, 53.99, and 77.38% when cell densities were 1081, 1176, and 1230 kg/m^3^, indicating that higher percentage of cells with the density of 1230 kg/m^3^ had more DNA contents.

**Figure 5.**
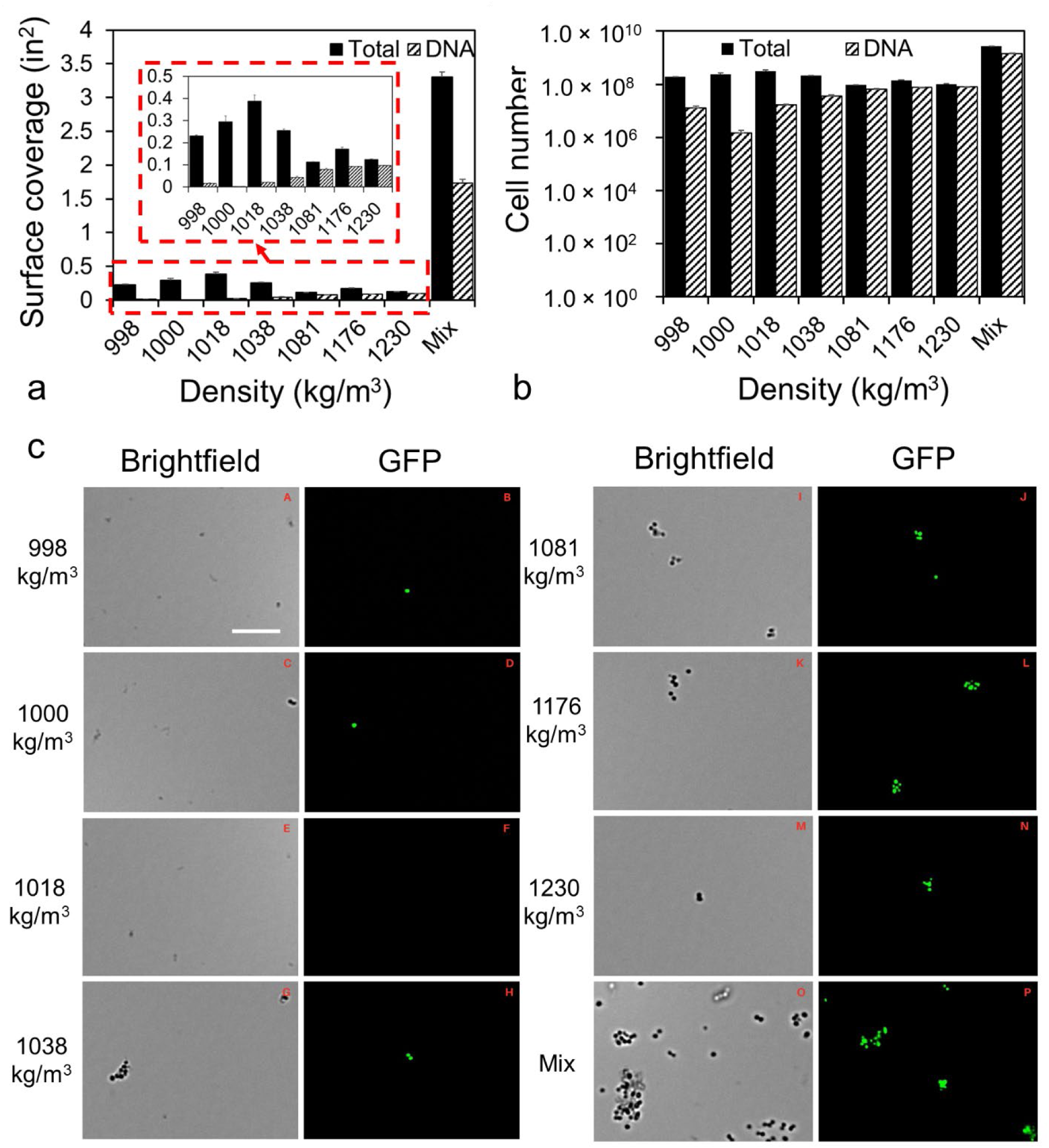
DNA content of cells with varying densities. Surface coverage (a) and the number (b) of cells with/without DNA labeled with green fluorescence. Surface coverage of total cell was characterized by analyzing the images taken by Brightfield Illumination. Surface coverage DNA was characterized by analyzing the images taken by green fluorescence [excitation: BP 470/40, FT 495 (HE), and BP 525/50 (HE)]. Five random images were taken for each condition as technical repeats and each condition was repeated biologically three times (N = 3). Standard errors are shown. Image analysis comparison of Brightfield and green fluorescence (c) Zoomed in images of Brightfield and green fluorescence channels for *S. aureus* cells with the density of 998, 1000, 1018, 1038, 1081, 1176, and 1230 kg/m^3^, respectively. The zoomed images of mixed *S. aureus* cells before separation are included as controls (Bar = 10 μm).

### 2.6 Cells with higher buoyant density were more susceptible to antibiotics

By directly visualizing the cells using a microscope, we found that cells with a higher buoyant density had more DNA content than those with lower buoyant density. To correlate this with the level of biofilm-related antibiotic resistance, we also quantified the level of antibiotic susceptibility of cells with various densities. As shown in Fig. 6, cells with a density of 1230 kg/m^3^ were more susceptible to the one-hour treatment with 100 μg/mL in 0.85% NaCl compared to cells with the densities of 998 and 1000 kg/m^3^ (t.test, *p* = 0.34). This difference in antibiotic susceptibility was solely attributed to cells with different densities because we matched sucrose background before the antibiotic treatment.

**Figure 6.**
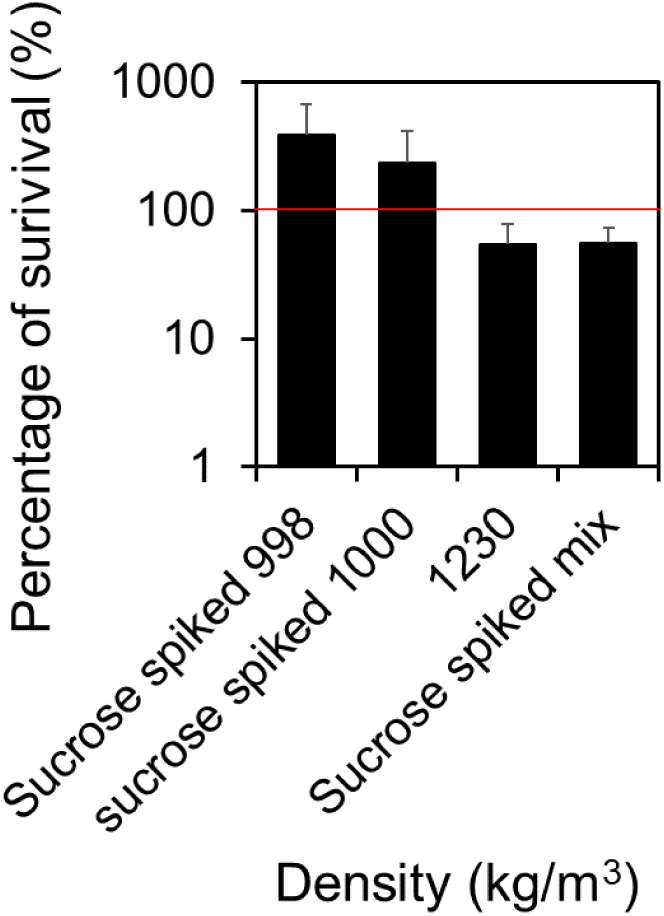
Susceptibility of cells with varying densities. The susceptibility of cells with the densities of 998, 1000, and 1230 kg/m^3^ were characterized by treating the cells separated using sucrose density gradient centrifugation with 100 μg/mL vancomycin for 1 hour in 0.85% NaCl at 37°C with shaking at 200 rpm. The number of cells survived the treatment was quantified using the most probable number (MPN) assay^32^. All experiments included in Fig. 6 were repeated biologically five times (N = 5). Standard errors are shown.

## 3. Discussion

*S. aureus* biofilms rank among the primary culprits for infectious diseases, encompassing conditions from skin afflictions to septicemia^33–35^. The treatment of biofilm-driven infections is challenging due to the restricted permeability of antimicrobial agents through the extracellular polymeric substances (EPS), coupled with the prevalence of cells exhibiting reduced growth and metabolic rates, a result of stress responses in mature biofilms^4, 5, 36, 37^. Despite the profound implications of biofilms, the complexity of their formation and emergent antibiotic resistance remains not fully deciphered. ^3, 8, 33, 38^. Biofilm evolution encompasses four key phases: initial adherence, microcolony creation, maturation, and dispersion^28, 39^. The initial attachment is critical to biofilm formation, but the exact factors dictating the shift from a planktonic state to a sessile one is yet to be entirely pinpointed. Notably, in contrast to bacteria *Escherichia coli* and *Pseudomonas aeruginosa*, *S. aureus* cells lack external structures, such as flagella or pili, to facilitate substrate engagement. The determinants governing the transition of *S. aureus* cells, particularly under forces like gravity and shear stress, remain veiled, thus sparking the interest behind this research.

Previously, it was well accepted that the density of bacterial cells, such as *S. aureus,* is closely resembled to that of water (998 kg/m^3^), when determining biomass. However, this assumption might be flawed. If accurate, *S. aureus* cells wouldn’t gravitate towards a substrate’s surface to initiate attachment, given the equilibrium between the cell’s weight and the buoyant force within the solution. Our gravity-driven sedimentation experiment revealed differing sedimentation rates amongst *S. aureus* cells within an isogenic population (Fig. 1). This disparity can primarily be linked to variations in the cells’ buoyant densities, indicating a heterogeneity, even within an isogenic *S. aureus* population.

This heterogeneity in *S. aureus* cells’ buoyant density was validated using sucrose density gradient centrifugation (Fig. 2). This phenomenon was consistent across an isogenic *S. aureus* population at various growth stages, though the cell distribution pattern fluctuated based on the growth phase. For instance, in an exponential phase of the *S. aureus* population, around 30.23 ± 7.59 and 31.74 ± 9.60 % cells exhibited mass densities of 1176 and 1230 kg/m^3^, respectively. Although the cell distribution during this phase might approximate a normal distribution, it would lean towards a broader curve. Previously, Kubitschek *et al.* reported that exponential phase *E. coli* cells adhered to a tighter normal distribution^10^. The distinct sedimentation behavior of *S. aureus* and *E. coli* can be attributed to differences in their external structures, like flagella and pili. This pattern evolved as, in a 15.5-hour old *S. aureus* sample, the percentages of cells exhibited mass densities of 1176 and 1230 kg/m^3^ dropped to 4.88 ± 2.58 and 4.68 ± 2.03% cells, respectively, and over 86.32% of cells demonstrated a density surpassing 1230 kg/m^3^. However, in an 18-hour-old sample, all *S. aureus* cells maintained a density at or below 1230 kg/m^3^, with the cell distribution deviating from the typical normal distribution, suggesting that the heterogeneity in cells’ buoyant density varies dynamically.

Given the density of the LB broth was measured as 980 kg/m^3^, the observed variation in density can be translated to fluctuations in buoyant density when samples were introduced into the LB broth for biofilm formation (with buoyant densities of *B* = −18, 20, 38, 58, 101, 196, or 250 kg/m^3^). Based on the correlation between sedimentation rate and buoyant density (Equation 1), we hypothesized that cells possessing a higher buoyant density would gravitate towards the substrate and initiate attachment sooner. This conjecture was subsequently validated through our biofilm assay (Fig. 3).

To investigate the mechanism behind the heterogeneity among cells in an isogenic *S. aureus* population, we analyzed the cellular content, particularly DNA, and growth rate of cells exhibiting different densities (Fig. 4&5). A greater content within a specific volume signifies a higher cell density (Fig. 5). Through microscopic examination of both the entire cell population and their DNA, we observed that subpopulations with densities of 998, 1000, 1018, and 1038 kg/m^3^ contained more cells than quantified via CFU. Intriguingly, the majority of these cells lacked DNA signals. Previously, Kim *et al*.^16^ found that viable but non-culturable (VBNC) and persisters often contain reduced or negligible cellular content. This leads to the inference that cells exhibiting low buoyant densities might either be VBNC or persisters. Cells with a low buoyant density, due to their minimal content, displayed fewer DNA signals compared to their high-density counterparts, suggesting that those cells might be VBNC or persisters. This is corroborated by their slower growth rate after cell reviving.

In evaluating the susceptibility of separated *S. aureus* subpopulations, we discerned that cells exhibiting a high buoyant density (1230 kg/m^3^) were notably more responsive to 100 μg/mL vancomycin than their lower buoyant density counterparts (998 and 1000 kg/m3) (Fig. 6). This mirrors the Gu *et a*l.’s prior findings^28^, wherein initially attached *E. coli* cells displayed an enhanced susceptibility to antibiotics because of higher level metabolic activities. However, these initially attached cells rapidly lose their antibiotic susceptibility due to matrix secretion and subsequent microcolony development. This transient heightened vulnerability in the initial phases may offer a unique opportunity for the effective elimination of *S. aureus* biofilms before they mature.

## 4. Conclusions

In this study, we found a heterogeneity in the buoyant density of isogenic *S. aureus* populations. This heterogeneity persists regardless of population’s age, but the cell distribution pattern evolves as the population ages, deviating from a narrow normal distribution. A higher buoyant density in *S. aureus* cells may be attributed to an increased cellular content, such as DNA. Intriguingly, cells demonstrating a higher buoyant density were more vulnerable to vancomycin, implying enhanced metabolic activity. It was supported by the 24-hour LB broth culture, that *S. aureus* cells with higher buoyant density more effectively formed biofilms on the 96-well plate. Our findings corroborate the hypothesis that cells with higher buoyant density, due to their intensified metabolic activities, gravitate towards and initially adhere to upright substrates. This pronounced susceptibility to antibiotics among the early-attaching cells presents a promising avenue for proactive biofilm management and eradication.

## 5. Materials and methods

### 5.1 Strains and growth media

*S. aureus ALC2085* was first streaked onto Lysogeny Broth (LB) Agar Plates that are made of 10 g/L of NaCl, 10 g/L tryptone, 5 g/L yeast extract, and 15 g/L agar. The plates were incubated at 37°C overnight. A colony was picked from the LB agar plate and used to inoculate fresh LB solution, to grow *S. aureus* cells at 37°C with shaking at 200 rpm.

### 5.2 Density gradient centrifugation separation

A series of sucrose solutions were prepared with diverse concentrations by dissolving sucrose (Sigma-Aldrich, St. Louis, MO) into 0.85% NaCl solutions, resulting in distinct mass densities: 998 (0.85% NaCl solution), 1000 (5% wt/vol sucrose), 1018 kg/m³ (10% wt/vol sucrose), 1038 kg/m³ (20% wt/vol sucrose), 1081 kg/m³ (30% wt/vol sucrose), 1176 kg/m³ (40% wt/vol sucrose), and 1230 kg/m³ (50% wt/vol sucrose). All sucrose solutions and bacterial cultures were chilled on ice for no less than 15 min before sucrose gradient centrifugation. Then, 100 µL of chilled bacterial culture was slowly aspirated on top of each solution. The gradient solutions in microcentrifuge tubes were centrifuged at 4°C and 170 × *g* for the first 5 minutes, and then 1,250 × *g* for another 5 minutes. After centrifugation, 100 µL from each layer of solutions were collected and placed in a 96-well plate.

Cell density in each layer of solutions was quantified using the drop plate method and colony forming units (CFU)^29^. In details, the 100 µL solutions from each layer were treated as the original samples and placed in the first column of the 96-well plate. These solutions were then diluted 10 times by transferring 20 μL of bacterial solution into 180 μL clean 0.85% NaCl for a dilution factor of 10. This step will be repeated n times to reach a dilution factor of 10^n^ in column n. Upon mixing, 10 μL from each well was dropped onto an LB agar plate. The excessive 0.85% NaCl in each 10 μL droplet was air dried to fix the bacteria onto the LB agar plate. The dried LB agar plates were incubated at 37°C overnight before the number of CFU can be counted.

### 5.3 Gravity-driven sedimentation

To corroborate the results obtained from density gradient centrifugation and prove that *S. aureus* populations are mixtures of cells with different buoyant density, we conducted a gravity-driven sediment. This assay was performed using test tubes filled with 0.85% NaCl solutions. Both bacterial cultures, test tubes, and 0.85% NaCl solution were pre-chilled on ice for no less than 15 min. In each test tube, 5 mL of 0.85% NaCl solution was added. Then, 500 μL of pre-chilled stationary phase *S. aureus* culture was carefully aspirated to the top of 5 mL cold 0.85% NaCl solutions in 15 mL Falcon Tubes. After 15 minutes, the top 500 µL was removed and the 0.85% NaCl solution was removed in 1 mL increments from the top. Cell number in each layer was quantified using drop plating and CFU as described above.

### 5.4 Biofilm inoculation

Cells from each layer separated using density gradient centrifugation above and cells before separation (mix) were first spiked with sucrose to 50% to match the sucrose concentration of the layer with the mass density of 1230 kg/m^3^. Then, cells were diluted using 96-well plates by following the method of dilution described above for drop plating. The difference is that instead of using 0.85% NaCl solutions for dilution, we used LB solution. After dilution, the number of cells in each well were quantified by dropping 10 µL cells onto LB agar plates and CFU. The 96-well plates were inoculated for 48 hours to form biofilms. After 48-h biofilm formation, a crystal violet assay^30^ was conducted. In detail, planktonic cells were dumped first and then followed with three washes using 0.85% NaCl solutions. Then, add 250 µL 0.1% solution of crystal violet in water into each well. Incubate the 96-well plate at room temperature for 10-15 min. Dump the dye and wash the stained biofilms three times using clean 0.85% NaCl solutions. Dry the 96-well plate overnight at room temperature by turning it upside down. The stained biofilms in each well were dissolved in 30% acetic acid in water. Absorbances were collected at 550 nm using a microplate reader (RayBiotech, Peachtree Corners, Georgia, United States) after 10-15 min incubation at room temperature. Guided with the CFU, only the biofilms initiated with similar cell numbers were compared.

### 5.5 Antibiotic susceptibility and most probable number (MPN) assay

Antibiotic susceptibility of *S. aureus* cells with different buoyant densities was evaluated by treating cells with 100 µg/mL vancomycin in 0.85% NaCl solutions for 1 hour at 37°C while shaking at 200 rpm. The 0.85% NaCl solutions were used to avoid cell duplications during treatment. Cells from each layer separated using density gradient centrifugation above and cells before separation (mix) were first spiked with sucrose to 50% to match the sucrose concentration of the layer with the mass density of 1230 kg/m^3^. Then, they were diluted in 0.85% NaCl solutions and LB solutions as described above for drop plating. Cells in each well were quantified using CFU. Cells diluted in LB solutions were directly incubated overnight at 37°C while shaking at 200 rpm to get the MNP^32^ of the control. Cells diluted in 0.85% NaCl solutions were treated with 100 µg/mL vancomycin for 1 hour at 37°C while shaking at 200 rpm. Treated samples were then transferred into a new 96 well plate, leaving a blank row of wells on each side of the well plate. Samples were diluted up to 10^9^ times with a dilution factor of 10 by transferring 20 μL of bacterial solution into 180 μL LB. The 96-well plates were then incubated at 37°C while shaking at 200 rpm. The number of cells after treatment was determined by using the MPN method^32^. In details, for each control or antibiotic treated sample, we treated the last dilution (e.g., 10^n^) that had change in OD_600_ compared to the LB control started with 1 cell. By doing this, we assumed that the original control or antibiotic treated sample had 10^n^ cells, regardless of the changes in the original antibiotic treated samples or the 10^1^ times dilution of the antibiotic treated samples. Due to the continuous treatment by 100 and 10 μg/mL vancomycin, during the growth in the original antibiotic treated samples or the 10^1^ times dilution of the antibiotic treated samples, there were no changes in the OD_600_. However, with the increase in the times of dilution (e.g., 10^n^), the concentration of vancomycin decreased to 100/10^n^ μg/mL that was too low to affect the growth of *S. aureus*. This was the reason why we could calculate back how many cells stayed viable using MNP after antibiotic treatment. Since the amount of viable cell number before treatment was also determined using MNP corroborated with CFU, we are confident that antibiotic susceptibility can be quantified with the presence of antibiotics.

### 5.6 DNA level quantification using Acridine Orange (AO) and fluorescence microscope

To quantify the DNA content of cells with different buoyant densities, we used Acridine orange (AO), a nucleic acid-selective fluorescent dye that can label DNA with green fluorescence (Ex. 500 nm and Em. 526 nm)^31^. In details, we prepared 5 mg/mL acridine orange (Sigma-Aldrich, St. Louis, MO) solution in DI water. Then, we diluted this working stock solution 10 times into the samples and incubated at room temperature for 3 min. After incubation, cells were imaged using a fluorescence microscope (Echo Revolve, BICO, Gothenburg, Sweden). Cells were illuminated with Brightfield (black vs. white contrast for all bacteria) and light source for exciting green fluorescence (for cells with DNA labeled with green fluorescence). Five images were taken at random spots for each condition. Each condition was repeated biologically three times.

Software Image J was used for quantifying the surface coverage (in^2^) of total cells and cells with DNA labeled with green fluorescence. For black and white images taken using Brightfield, the images are uploaded, and the threshold was set individually to achieve a clear contrast between the background and the bacteria. The bacteria should be displayed as small white particles shown clearly on a black background. For images of cells with green fluorescence, images were altered to be 16 bits first and then the threshold was set the same way as described prior for Brightfield. By doing this, background was removed and only bacteria can be seen in the images. The areas of surface coverage were calculated using the Analyze Particle’s Function in Image J.

### 5.7 Statistics

For density gradient centrifugation, antibiotic susceptibility test, and biofilm formation at air-liquid and liquid-solid interfaces, each experiment will be conducted with at least three biological replicates (n ≥ 3) based on a power analysis (n = 0.34 < 3 when confidence α = 0.05 and power 0.90) and studies reported previously (n ≥ 3)^28, 40^. For tracking persisters during bacterial growth using microfluidic devices and microcapsules, at least three biological replicates (n ≥ 3) and 500 bacteria from each biological replicate will be analyzed. For fluorescence imaging and tracking, at least five technical repeats will be taken from random spots during each biological replication, and ∼120 bacteria will be tracked and analyzed for each repeat^41^. Statistical design of experiments, such as power analysis, factorial design, and complete randomization block design, will be applied as appropriate. Collected data will be analyzed using a *t*-test, *z*-test, correlation coefficient, Analysis of Variance (ANOVA), and Bonferroni method as appropriate.

## 6. Acknowledgements

The author thanks the NASA CT Space Grant (80NSSC20M0129) for supporting this research.

## 7. Author contributions

H. Gu conceived the idea. H. Gu, M. Liguori, and S. Kispert designed the experiments. M. Liguori, S. Kispert, and C. Valikaneye conducted the experiments. C. Qiu, T. H.R. Niepa, S. Wang, and N. Zhang participated in the manuscript preparation.

